# The medial orbitofrontal cortex - basolateral amygdala circuit regulates the influence of reward cues on adaptive behavior and choice

**DOI:** 10.1101/2021.04.27.441665

**Authors:** Nina T. Lichtenberg, Linnea Sepe-Forrest, Zachary T. Pennington, Alexander C. Lamparelli, Venuz Y. Greenfield, Kate M. Wassum

**Author notes:** Correspondence: Kate Wassum, Dept. of Psychology, UCLA, 1285 Pritzker Hall, Box 951563, Los Angeles, CA 90095-1563.

## Abstract

Adaptive reward-related decision making requires accurate prospective consideration of the specific outcome of each option and its current desirability. Often this information must be inferred based on the presence of predictive environmental events. The basolateral amygdala (BLA) and medial orbitofrontal cortex (mOFC) are two key nodes in the circuitry supporting such outcome expectations, but very little is known about the function of direct connections between these regions. Here, in male rats, we first anatomically confirmed the existence of bidirectional, direct projections between the mOFC and BLA and found that BLA projections to mOFC are largely distinct from those to lateral OFC (lOFC). Next, using pathway-specific chemogenetic inhibition and the outcome-selective Pavlovian-to-instrumental transfer and devaluation tests, we interrogated the function of the bidirectional mOFC-BLA connections in reward-directed behavior. We found evidence that the mOFC→BLA pathway mediates the use of environmental cues to understand which specific reward is predicted, information needed to infer which action to choose, and how desirable that reward is to ensure adaptive responses to the cue. By contrast, the BLA→mOFC pathway is not needed to use the identity of an expected reward to guide choice, but does mediate adaptive responses to cues based on the current desirability of the reward they predict. These functions differ from those we previously identified for the lOFC-BLA circuit. Collectively, the data reveal the mOFC-BLA circuit as critical for the cue-dependent reward outcome expectations that influence adaptive behavior and decision making.

**SIGNIFICANCE STATEMENT:** To make good decisions we evaluate how advantageous a particular course of action would be. This requires understanding what rewarding outcomes can be expected and how desirable they currently are. Such prospective considerations are critical for adaptive decision making but disrupted in many psychiatric diseases. Here we reveal that direct connections between the medial orbitofrontal cortex and basolateral amygdala mediate these functions. These findings are especially important in light of evidence of dysfunction in this circuit in substance use disorder and mental illnesses marked by poor decision making.

---

To make good decisions we evaluate how advantageous a particular course of action would be, given our circumstances, or state. This includes consideration of what outcomes (e.g., specific rewarding events) might occur as well as the current desirability of those potential outcomes. Often the actual outcome we could receive is not readily apparent in the immediate environment. In these cases, we infer the most advantageous option by mentally representing possible outcomes and their current values. These representations are facilitated by predictive environmental events. Stored stimulus-outcome and action-outcome associative memories enable the outcome expectations that influence decision making (Balleine and Dickinson, 1998; Delamater, 2012; Fanselow and Wassum, 2015). For example, a restaurant logo on your favorite food-delivery app lets you know that a specific type of food (e.g., tacos) is available and, based on your current state (e.g., did you just have Mexican for lunch?), you can determine that food's value and decide if it is a suitable dinner option. Such prospective considerations are critical for adaptive decision making and disrupted in many psychiatric diseases, yet, much is unknown of their underlying neural circuits.

The orbitofrontal cortex (OFC) and basolateral amygdala (BLA) are two key nodes in the circuitry supporting reward outcome representations and their influence over decision making. Damage to either region causes performance deficits when adaptive behavior requires one to represent possible rewarding events (Hatfield et al., 1996; Gallagher et al., 1999; Blundell et al., 2001; Pickens et al., 2003; Izquierdo et al., 2004; Corbit and Balleine, 2005; Pickens et al., 2005; Wellman et al., 2005; Machado and Bachevalier, 2007; Ostlund and Balleine, 2007, 2008; Johnson et al., 2009; West et al., 2011; Rhodes and Murray, 2013; Malvaez et al., 2015; Lichtenberg and Wassum, 2016; Panayi and Killcross, 2018). The lateral OFC (lOFC) has been particularly implicated, but the anatomically and functionally distinct medial OFC (mOFC) (Wallis, 2011; Izquierdo, 2017; Woon et al., 2020) also participates in appetitive decision making (Noonan et al., 2010; Stopper et al., 2014; Bradfield et al., 2015; Dalton et al., 2016; Noonan et al., 2017; Bradfield et al., 2018; Malvaez et al., 2018a; Münster and Hauber, 2018), especially when the value of rewarding options must be mentally simulated (Bray et al., 2010; Bradfield et al., 2015) and/or compared (Noonan et al., 2010; Rudebeck and Murray, 2011; Stopper et al., 2014; Gourley et al., 2016; Yamada et al., 2018). The mOFC both sends and receives direct, excitatory projections to and from the BLA, positioning this as a potential crucial circuit for the influence of reward representations over decision making and adaptive behavior. But little is known of the function of direct mOFC→BLA projections and nothing of BLA→mOFC projections. This is limiting for both basic understanding of appetitive behavior as well as for the understanding and treatment of the myriad psychiatric illnesses marked by dysfunction in the mOFC-BLA circuit (Tanabe et al., 2009; Goldstein and Volkow, 2011; Hahn et al., 2011; Linke et al., 2012; Sladky et al., 2015; Xie et al., 2021).

To address this gap in knowledge, we used a pathway-specific chemogenetic inhibition approach in male rats to uncover the function of mOFC→BLA and BLA→mOFC projections in the prospective consideration of specific predicted rewards and their current desirability required for adaptive reward-related behavior. We used Pavlovian and instrumental conditioning procedures that engender the encoding of two distinct, appetitive, sensory-specific stimulus-outcome memories and two distinct action-outcome memories. We followed this with an outcome-specific Pavlovian-to-instrumental transfer (PIT) test, which assesses the use of these memories to allow reward cues to influence decision making, and an outcome-selective devaluation test to evaluate the ability to use the value of an anticipated reward to guide adaptive behavior. Procedures were identical to our prior investigation of the function of the lOFC-BLA circuit (Lichtenberg et al., 2017) to facilitate comparison of function of the pathways between the BLA and lOFC v. mOFC.

## MATERIALS AND METHODS

### Subjects

Male, Long Evans rats aged 10-12 weeks at the start of the experiment (Charles River Laboratories, Wilmington, MA) were pair housed prior to surgery and then subsequently housed individually in a temperature (68-79°F) and humidity (30-70%) regulated vivarium. Rats were provided with filtered tap water *ad libitum* in the home cage and were maintained on a food-restricted 12-14 g daily diet (Lab Diet, St. Louis, MO) to maintain ~85-90% free-feeding body weight. Rats were handled for ~3 days prior to experiment onset. Separate groups of naïve rats were used for each experiment. Experiments were performed during the dark phase of a 12:12 hr reverse dark/light cycle (lights off at 7AM). All procedures were conducted in accordance with the NIH Guide for the Care and Use of Laboratory Animals and were approved by the UCLA Institutional Animal Care and Use Committee.

### Surgery

Standard surgical procedures, described previously (Malvaez et al., 2015; Lichtenberg et al., 2017; Malvaez et al., 2019), were used for all surgeries. Rats were anesthetized with isoflurane (4–5% induction, 1–2% maintenance) and a nonsteroidal anti-inflammatory agent was administered pre- and post-operatively to minimize pain and discomfort. Rats also received post-operative antibiotics.

#### Anatomical tracing of mOFC-BLA circuit

Rats (*N* = 3) were infused unilaterally into the BLA with an anterograde and retrograde viral tracer cocktail. The anterograde adeno-associated virus (AAV) expressing the mCherry fluorophore (AAV8-hsyn-mCherry, 4.6 × 10^12^ vg/mL; University of North Carolina Vector Core, Chapel Hill, NC) was mixed 1:1 with the retrogradely trafficked AAV encoding eGFP (AAVrg-hsyn-EGFP, 7.4 × 10^12^ vg/mL; Addgene, Watertown, MA) and infused into the BLA (AP: −3.0, ML: ±5.1, DV: −8.6 mm from bregma; 0.5 μL) at a rate of 0.1 μL/min using a 33-gauge injector. Injectors were left in place for an additional 10 minutes to ensure adequate diffusion and to minimize off-target spread along the injector tract. Tissue was collected 8 weeks post-surgery to ensure anterograde and retrograde transport for robust expression in BLA axons and terminals in the mOFC and BLA-projecting mOFC cell bodies.

#### Anatomical tracing of BLA projections to mOFC and lOFC

Rats (*N* = 4) were infused unilaterally with AlexaFluor-594 conjugated cholera toxin B (CTb; 5 μg/μL; Life Technologies, San Diego, CA) into the mOFC (AP +4.1, ML ±0.7, DV −5.0 mm from bregma; 0.4 μL) and AlexaFluor-488 conjugated CTb into the lOFC (AP +3.0, ML ±3.2, DV −5.8 mm; 0.6 μL) at a rate of 0.1 μL/min using a 33-gauge injector. Injectors were left in place for an additional 10 minutes. Tissue was collected 2 weeks post-surgery.

#### Chemogenetic inhibition of mOFC→BLA projections

Surgery occurred prior to onset of behavioral training. Rats (final *N* = 11) were infused bilaterally with AAV expressing the inhibitory designer receptor *human M4 muscarinic receptor* (hM4Di; AAV8-hSyn-hM4Di-mCherry, 4.8 × 10^12^ or 3.7 × 10^12^ vg/mL; Addgene; 0.3 μL) at a rate of 0.1 μL/min into the mOFC (AP: +4.1, ML: ±0.5, DV: −5.2 mm from bregma) using a 33-gauge injector. Injectors were left in place for an additional 10 minutes to ensure adequate diffusion and to minimize off-target spread along the injector tract. 22-gauge stainless-steel guide cannulas were implanted bilaterally above the BLA (AP: −3.0, ML: ±5.1, DV: −7.0 mm from bregma). Rats recovered for ~16 days before the onset of behavioral training. Behavioral testing began 6-7 weeks following surgery to ensure anterograde transport and robust expression in mOFC axons and terminals in the BLA. 1 rat from this group became ill during behavioral training and was humanely euthanized before testing.

#### Chemogenetic inhibition of BLA→mOFC projections

Surgery occurred prior to onset of behavioral training. Rats (final *N* = 9) were infused bilaterally with AAV expressing the inhibitory designer receptor hM4Di (AAV8-hSyn-hM4Di-mCherry, 4.8 × 10^12^ or 3.7 × 10^12^ vg/mL; Addgene; 0.4 μL) at a rate of 0.1 μL/min into the BLA (AP: −3.0, ML: ±5.1, DV: −8.6 mm from bregma) using a 33-gauge injector. Injectors were left in place for an additional 10 minutes. A 23-gauge stainless-steel guide cannula was implanted bilaterally above the mOFC (AP: +4.1, ML: ±0.7, DV: −2.8 mm from dura). Rats recovered for ~16 days before the onset of behavioral training. Behavioral testing began 6-7 weeks following surgery to ensure anterograde transport and robust expression in BLA axons and terminals in the mOFC.

### Behavioral Procedures

#### Apparatus

Training took place in Med Associates conditioning chambers (East Fairfield, VT) housed within sound- and light-attenuating boxes, described previously (Malvaez et al., 2015; Lichtenberg and Wassum, 2016; Wassum et al., 2016; Lichtenberg et al., 2017; Collins et al., 2019; Malvaez et al., 2019). Each chamber contained 2 retractable levers that could be inserted to the left and right of a recessed food-delivery port in the front wall. A photobeam entry detector was positioned at the entry to the food port. Each chamber was equipped with a syringe pump to deliver 20% sucrose solution in 0.1 mL increments through a stainless-steel tube into one well of the food port and a pellet dispenser to deliver single 45-mg grain food pellets (Bio-Serv, Frenchtown, NJ) into another well of the same food port. Both a tone and white noise generator were attached to individual speakers on the wall opposite the levers and food port. A 3-watt, 24-volt house light mounted on the top of the back wall opposite the food-delivery port provided illumination and a fan mounted to the outer chamber provided ventilation and external noise reduction. Behavioral procedures were similar to those we described previously (Malvaez et al., 2015; Lichtenberg and Wassum, 2016; Lichtenberg et al., 2017).

#### Pavlovian conditioning

Rats first received 8 sessions of Pavlovian training (1 session/day) to learn to associate each of two auditory conditional stimuli (CSs; 80-82 db, 2-min duration), tone (1.5 kHz) or white noise, with a specific food reward, sucrose (20%, 0.1 mL/delivery) or grain pellets (1 45 mg pellet/delivery; Bio-Serv). CS-reward pairings were counterbalanced at the start of each experiment. For half the subjects, tone was paired with sucrose and noise with pellets, with the other half receiving the opposite arrangement. Each session consisted of 8 tone and 8 white noise presentations. During each 2-min CS the associated reward was delivered on a 30-s random-time schedule, resulting in an average of 4 stimulus-reward pairings per trial. CSs were delivered in pseudorandom order with a variable 2 - 4 min (mean = 3 min) intertrial interval.

#### Instrumental conditioning

Rats were then given 11 days, minimum, of instrumental training. They received 2 separate training sessions per day, one with the left lever and one with the right lever, separated by at least 1 hr. Each action was reinforced with a different outcome (e.g., left press-pellets / right press-sucrose; counterbalanced with respect to the Pavlovian contingencies). Each session terminated after 30 outcomes had been earned or 30 min had elapsed. The reinforcement schedule was escalated ultimately to random-ratio 20 (RR-20; on average 20 presses required to obtain reward delivery). Rats received one training day (minimum) in which their actions were continuously reinforced, then a minimum of 2 training days on RR-2, 2 days on RR-5, and 3 days on RR-10, before being moved to the final RR-20 schedule for at least 3 days. To be escalated to the next schedule, rats had to earn at least 24 outcomes within 30 min. This requirement had to be met on the final RR-20 schedule for at least 2 training days before subjects were advanced to testing. The maximum number of days required to reach criterion for instrumental acquisition was 15.

#### Outcome-selective Pavlovian-to-instrumental transfer test

Following Pavlovian and instrumental conditioning, rats received a set of outcome-selective Pavlovian-to-instrumental transfer (PIT) tests. On the day prior to each PIT test, rats were given a single 30-min extinction session during which both levers were available but pressing was not reinforced to establish a low level of responding. During the PIT test, both levers were continuously present, but pressing was not reinforced. After 5 min of lever-pressing extinction, each 2-min CS was presented separately 4 times in pseudorandom order, separated by a fixed 4-min intertrial interval. No rewards were delivered during CS presentation. Rats received two of each test, one following infusion of vehicle and one following infusion of CNO, counterbalanced for order. Rats were given 2 retraining sessions for each instrumental association (2 sessions/day for 2 days) and 1 Pavlovian retraining session in between tests.

#### Outcome-selective devaluation test

Rats were given a set of 2 outcome-specific devaluation tests. Immediately before each test, rats were given 1-hr, unlimited access to either sucrose solution or food pellets in pre-exposed feeding chambers to establish a sensory-specific satiety and thus selectively devalue the prefed reward. The test consisted of two phases. In the first, both levers were available, and nonreinforced lever pressing was assessed for 5 min. The levers were then retracted, which started the second, Pavlovian, test phase, in which each 2-min CS was presented, without accompanying reward, separately 2 times each in alternating order, separated by a fixed, 4-min intertrial interval. Rats received 2 of each test, one in which food pellets were devalued and one in which sucrose solution was devalued. After prefeeding and immediately prior to the test rats were infused with either vehicle or CNO. Test order and drug/prefed outcome relationship were counterbalanced across subjects and with respect to the CS-reward relationships. For both the mOFC→BLA and BLA→mOFC groups, there were no significant differences in the amount consumed during prefeeding between the future drug conditions (mOFC→BLA, Vehicle: 21.14 ± 2.60 s.e.m. grams; CNO: 22.35 ± 3.06; *t*_9_ = 0.24, *P* = 0.81; BLA→mOFC, Vehicle: 19.02 ± 2.28; CNO: 16.58 ± 2.96; *t*_8_ = 0.58, *P* = 0.57). Successful devaluation was confirmed by post-test consumption of each food reward. For both the mOFC→BLA (Vehicle, Valued: 8.26 ± 1.35 s.e.m. grams, Devalued: 0.92 ± 0.31; CNO, Valued: 7.45 ± 1.26, Devalued: 1.81 ± 0.78; Outcome (Valued v. Devalued): *F*(_1,9_) = 76.06, *P* < 0.0001; Drug: *F*(_1,9_) = 0.0009, *P* = 0.98; Drug × Outcome: *F*(_1,9_) = 0.47, *P* = 0.51) and BLA→mOFC (Vehicle, Valued: 4.81 ± 0.75, Devalued: 3.82 ± 1.24; CNO, Valued: 9.36 ± 2.07, Devalued: 1.49 ± 0.56; Outcome: *F*(_1,8_) = 17.86, *P* = 0.003; Drug: *F*(_1,8_) = 3.21, *P* = 0.11; Drug × Outcome: *F*(_1,8_) = 3.95, *P* = 0.08) groups, across both drug conditions rats ate significantly less of the devalued food than the valued food. We excluded 1 mOFC→BLA rat from the devaluation test data set because he did not eat during one of the prefeeding sessions. Rats were given 1-2 days without behavioral training or testing after each devaluation test to reestablish hunger and then received 2 retraining sessions for each instrumental association (2 sessions/day for 2 days) and 1 Pavlovian retraining session in between tests.

#### Data collection

Lever presses and/or discrete entries into the food-delivery port were recorded continuously for each session. For the Pavlovian training and test sessions, the 2-min periods prior to each CS onset (preCS) served as the baseline for comparison of CS-induced elevations in lever pressing and/or food-port entries.

### Chemogenetic inhibition of mOFC→BLA projections

In the mOFC→BLA group, chemogenetic inhibition was used to inactivate hM4Di-expressing mOFC axons and terminals in the BLA prior to one PIT test and after prefeeding prior to one of the devaluation tests. We selected chemogenetic inhibition to allow inhibition throughout the duration of each test. Clozapine-*n*-oxide (CNO; Tocris Bioscience, Sterling Heights, MI) was dissolved in artificial cerebral spinal fluid (aCSF) to 1 mM and 0.5 μL was intracranially infused over 1 min bilaterally into the BLA as previously described (Lichtenberg et al., 2017; Malvaez et al., 2019). Injectors were left in place for at least 1 additional min to allow for drug diffusion. Testing commenced within 5-10 min following infusion. Using these procedures, we have previously demonstrated effective inactivation of OFC axons and terminals in the BLA both *in vivo* and *ex vivo* (Lichtenberg et al., 2017; Malvaez et al., 2019). We have also demonstrated that this dose of CNO when infused into the BLA has no effect on reward-related behavior, BLA activity, or OFC terminal activity in the BLA in the absence of the hM4Di transgene (Lichtenberg et al., 2017; Malvaez et al., 2019). Thus, for the control condition aCSF was infused into the BLA using identical procedures.

### Chemogenetic inhibition of BLA→mOFC projections

For the BLA→mOFC group, chemogenetic inhibition was used to inactivate hM4Di-expressing BLA axons and terminals in the mOFC prior to one PIT test and after prefeeding prior to one of the devaluation tests. Procedures were identical to those above with the exception that CNO or aCSF was infused bilaterally into the mOFC (0.3 μL). We have previously shown these procedures to effectively attenuate the activity of BLA terminals in the OFC and have found that intra-OFC infusion of this dose of CNO does not affect reward-related behavior, OFC activity, or BLA terminal activity in the OFC in the absence of hM4Di (Lichtenberg et al., 2017). 3 rats were excluded from this group due to cannula clog and thus inability to infuse CNO.

### Histology

Rats were deeply anesthetized with Nembutal and transcardially perfused with phosphate buffered saline (PBS) followed by 4% paraformaldehyde (PFA). Brains were removed and post-fixed in 4% paraformaldehyde overnight, placed into 30% sucrose solution, then sectioned into 30-40 μm slices using a cryostat and stored in PBS or cryoprotectant. For the tracing experiments, fluorescent microscopy was used to confirm expression of eYFP in BLA-projecting mOFC cell bodies. Free-floating coronal sections were mounted onto slides and coverslipped with ProLong Gold mounting medium with DAPI (Invitrogen, Carlsbad, CA). The signal for BLA axonal expression of mCherry in the mOFC was immunohistochemically amplified using antibodies directed against mCherry. Floating coronal sections were washed 2 times in 1X PBS for 10 min and then blocked in a solution of 5% normal goal serum (NGS) and 1% Triton X-100 dissolved in PBS for 1-2 hrs at room temperature. Sections were then washed 3 times in PBS for 15 min and then incubated in blocking solution containing rabbit anti-DsRed antibody (1:1000; EMD Millipore, Billerica, MA) with gentle agitation at 4°C for 18-22 hrs. Florescence imaging was used to visualize AlexaFluor-488, and 594 conjugated CTb in lOFC- and mOFC-projecting BLA cells.

To visualize hM4Di-mCherry expression in BLA or mOFC cell bodies, free-floating coronal sections were mounted onto slides and coverslipped with ProLong Gold mounting medium with DAPI (Invitrogen). Axonal expression of hM4Di-mCherry in terminal regions was immunohistochemically amplified. Floating coronal sections were washed 2 times in 1X PBS for 10 min and then blocked in a solution of 5% NGS and 1% Triton X-100 dissolved in PBS for 1-2 hrs at room temperature. Sections were then washed 3 times in PBS for 15 min and then incubated in blocking solution containing rabbit anti-DsRed antibody (1:1000; EMD Millipore, Billerica, MA) with gentle agitation at 4°C for 18-22 hrs. On the second day, sections were rinsed 3 times in the blocking solution and incubated in Alexa Fluor 594-conjugated (red) goat secondary antibody (1:500; Invitrogen) for 2 hr.

Images were acquired using a Keyence BZ-X710 microscope (Keyence, El Segundo, CA) with a 4x,10x, and 20x objective (CFI Plan Apo), CCD camera, and BZ-X Analyze software. Subjects with off-target viral and/or cannula placements were removed from the dataset (mOFC→BLA: *N* = 7; BLA→mOFC: *N* = 7). One subject from the mOFC→BLA group had extensive tissue damage and was also removed. We also excluded *N* = 2 from the mOFC→BLA group and *N* = 1 from the BLA→mOFC group because we were unable to verify expression.

### Data analysis

#### Behavioral analysis

Behavioral data were processed with Microsoft Excel (Microsoft, Redmond, WA). Left and/or right lever presses and/or entries into the food-delivery port were collected continuously for each training and test session. For the last day of Pavlovian training, we compared the rate of food-port entries between the CS probe (after CS onset, before reward delivery) and the preCS baseline periods. Data were averaged across trials for each CS and then averaged across the two CSs. Press rates on the last day of instrumental training were averaged across levers. For the PIT test, baseline lever-press rate (presses/min) averaged across both levers during the 2-min periods immediately prior to the onset of each CS was compared with that during the CS periods. During the CS periods, lever pressing was separated for presses on the lever that, during training, earned the same outcome as the presented cue (CS-Same presses) versus those on the other available lever (CS-Different presses). Data were averaged across trials for each CS and then averaged across the two CSs. Rate of entries into the food-delivery port were also compared between the baseline and CS periods, averaged across both CSs. For the devaluation test instrumental phase, lever-press rate was compared between the lever that, in training, earned the prefed food outcome (i.e., the devalued outcome) and that which had previously earned the non-prefed (valued) outcome. During the Pavlovian phase the rate of entries into the food-delivery port were compared between the 2-min preCS baseline periods and the CS periods, which were separated by the CS that predicted the devalued outcome v. the CS that predicted the valued outcome.

#### Image Analysis

For the CTb tracing experiment, for each subject we analyzed 2 slices each of the anterior (−2.3 to −2.56 mm posterior to bregma), middle (−2.8 to −3.14 mm posterior to bregma), and posterior (−3.3 to −3.6) BLA. We used 20X images from the center of the lateral amygdala (LA) and BLA for each analysis. Single- and double-labeled neurons for each fluorophore were counted using a custom-written script in ImageJ (National Institutes of Health, Bethesda, MD, v. 1.50i). Cells counts were totaled across the LA and BLA and across slices.

#### Statistical analysis

Datasets were analyzed by two-tailed, paired Student's *t* tests, one- or two-way repeated-measures analysis of variance (ANOVA), as appropriate (GraphPad Prism, GraphPad, San Diego, CA; SPSS, IBM, Chicago, IL). Post hoc tests used the Bonferroni correction. All data were tested for normality prior to analysis with ANOVA and the Greenhouse-Geisser correction was applied to mitigate the influence of unequal variance between conditions if the assumption of sphericity was not met. Alpha levels were set at *P* < 0.05.

### Rigor and reproducibility

Group sizes were estimated *a priori* based on prior work using male Long Evans rats in this behavioral task (Malvaez et al., 2015; Lichtenberg and Wassum, 2016; Lichtenberg et al., 2017) and to ensure counterbalancing of CS-reward and Lever-reward pairings. Investigators were not blinded to condition because they were required to administer drug. All behaviors were scored using automated software (MedPC). Each behavioral experiment included at least 1 replication cohort and cohorts were balanced by viral group, CS-reward and Lever-reward pairings, Drug test order, etc. prior to the start of the experiment.

## RESULTS

### The BLA and mOFC are bidirectionally connected

Using anterograde and retrograde viral-mediated tracing, we first confirmed the existence of direct connections between the BLA and mOFC, as identified previously (Kita and Kitai, 1990; Hoover and Vertes, 2011; Heilbronner et al., 2016; Reppucci and Petrovich, 2016; Malvaez et al., 2019; Barreiros et al., 2021). An anterograde and retrograde tracer was infused into the BLA of male rats (Figure 1a; *N* = 3). We detected robust expression of the retrograde tracer fluorophore in BLA-projecting mOFC cell bodies, confirming a direct mOFC→BLA projection. We also detected robust axonal expression of the anterograde tracer fluorophore in the mOFC, indicating the presence of BLA→mOFC projections. Thus, the mOFC and BLA share direct, bidirectional connections.

**Figure 1.**
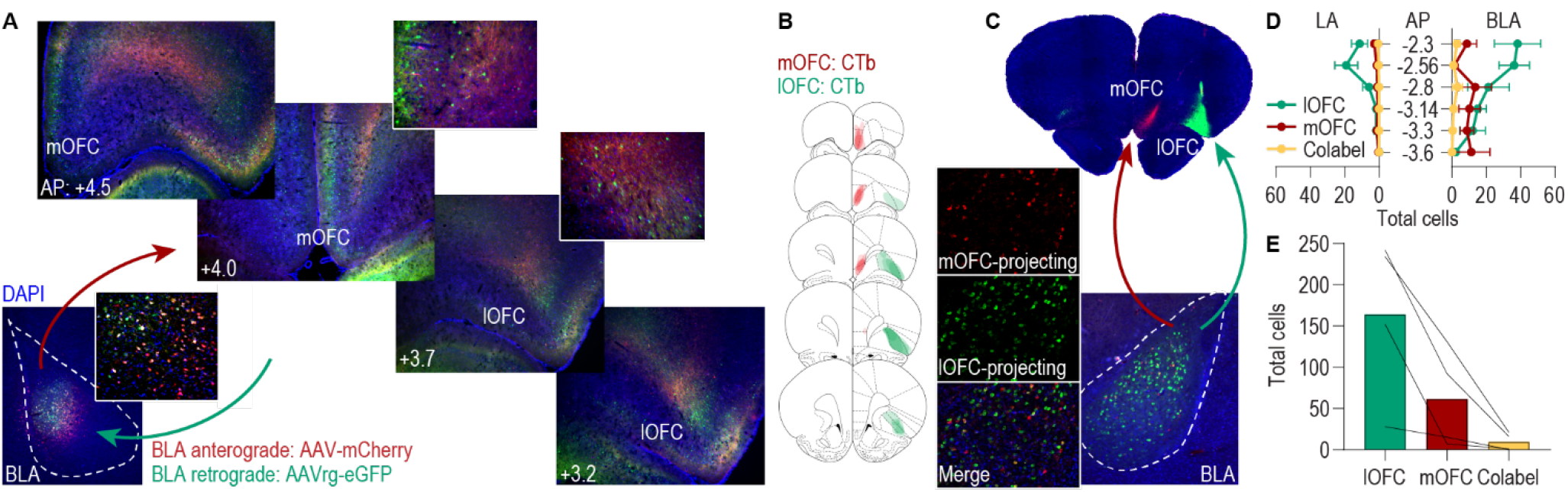
The BLA and mOFC are bidirectionally connected. **(a)** Representative images of the anterograde (mCherry) and retrograde (eGFP) tracer expression in the BLA infusion site (lower left) and of BLA axons and terminals (mCherry) and BLA-projecting cell bodies (eGFP) throughout the medial and lateral OFC. Insets show 20x image of cell body and axonal expression. **(b)** Schematic representation of the spread of the retrograde tracers AlexaFluor-594 CTb (red; mOFC) and AlexaFluor-488 CTb (green; lOFC) in the OFC infusion sites. **(c)** Representative images of AlexaFluor-594 CTb and AlexaFluor-488 CTb spread in the mOFC and lOFC infusion sides, and of mOFC- and lOFC-projecting cell bodies in the BLA. **(d)** Quantification of AlexaFluor-594 CTb (mOFC-projecting), AlexaFluor-488 CTb-labeled (lOFC-projecting), and co-labeled cells across the anterior-posterior (AP) axis of the lateral amygdala (LA) and basolateral amygdala (BLA). **(e)** Quantification of AlexaFluor-594 CTb (mOFC-projecting), AlexaFluor-488 CTb-labeled (lOFC-projecting), and co-labeled cells totaled across LA and BLA for all slices.

The BLA also projects to the lOFC (Morecraft et al., 1992; Lichtenberg et al., 2017; Barreiros et al., 2021) and we previously found that activity in these projections participates in decision making and adaptive behavior (Lichtenberg et al., 2017). Thus, we next asked whether the BLA pathway to the mOFC is distinct or overlapping with that to the lOFC. That is, whether the BLA sends collateral projections to both mOFC and lOFC. We injected retrograde tracers into the lOFC and mOFC of male rats (Figure 1b–c; *N* = 4). We detected expression of both tracer fluorophores in BLA cell bodies across the anterior-posterior (AP) extent of the BLA, confirming the presence of BLA projections to both lOFC and mOFC (Figure 1d; Lateral amygdala, AP × Projection: *F*(_10,30_) = 4.95, *P* = 0.0003; AP: *F*(_5,15_) = 4.10, *P* = 0.02; Projection: *F*(_2,6_) = 8.23, *P* = 0.02; BLA, AP × Projection: *F*(_10,30_) = 4.21, *P* = 0.001; AP: *F*(_5,15_) = 1.31, *P* = 0.31; Projection: *F*(_2,6_) = 8.78, *P* = 0.001). lOFC-projecting cells were more prominent than mOFC projectors in the anterior BLA. We detected more labeling of BLA→lOFC projecting cells than BLA→mOFC projecting cells, likely owing to the larger size, and thus tracer infusion volume, of the lOFC relative to mOFC. Critically, we detected very few co-labeled cells, indicating that, for the most part, the BLA pathways to the mOFC and lOFC are distinct (Figure 1e; *F*(_2,6_) = 10.11, *P* = 0.01).

### The mOFC→BLA pathway regulates the influence of stimulus-outcome memories over decision making and adaptive conditional responding

We next used chemogenetic inhibition to interrogate the function of mOFC→BLA projections in the influence of stimulus-outcome and action-outcome memories over decision making and adaptive conditional behavior (Figure 2a). We expressed the inhibitory designer receptor human M4 muscarinic receptor (hM4Di) in mOFC and placed guide cannula bilaterally over the BLA near hM4Di-expressing mOFC axons and terminals (Figure 2b–c; *N* = 11). This allowed us to later infuse clozapine-*n*-oxide (CNO; 1 mM) to inactivate mOFC axons and terminals in the BLA, as we have previously validated both *in vivo* and *ex vivo* (Lichtenberg et al., 2017; Malvaez et al., 2019).

**Figure 2.**
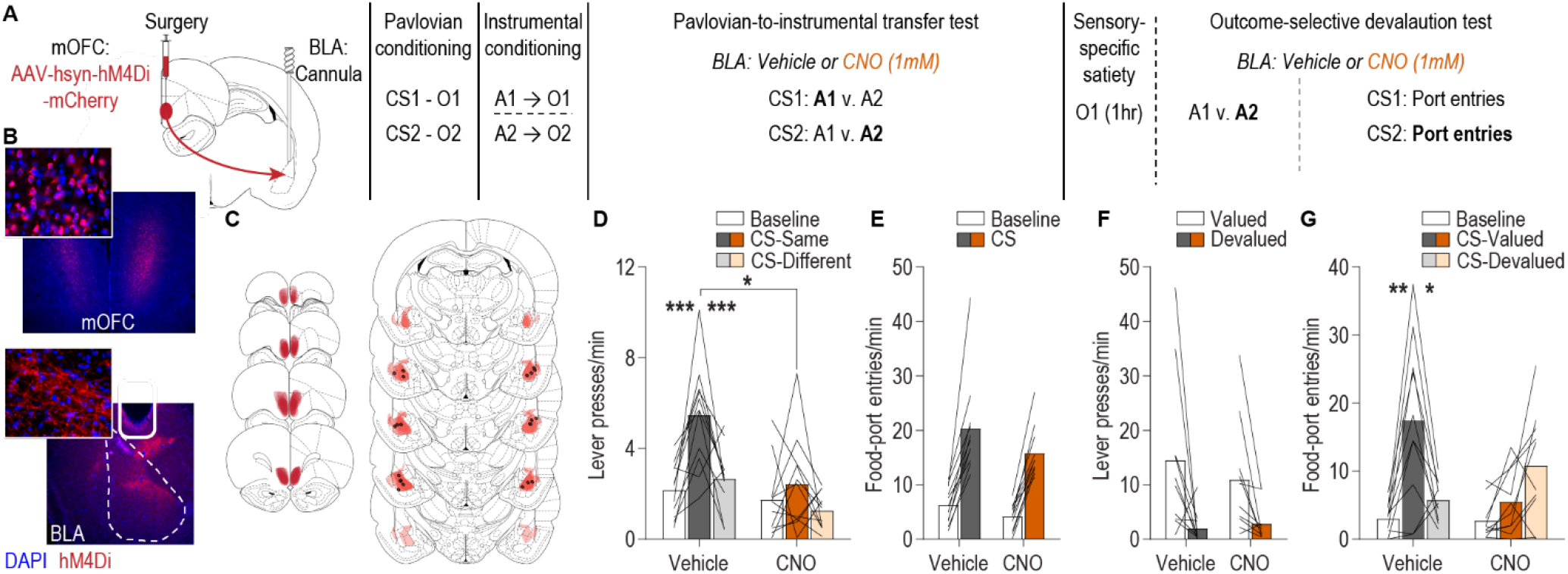
The mOFC→BLA pathway regulates the influence of stimulus-outcome memories over decision making and adaptive conditional responding. **(a)** Schematic of behavioral training, testing, and chemogenetic inactivation strategy. CS, conditional stimulus (white noise or tone); O, outcome (sucrose solution or food pellet); A, action (left or right lever press); CNO, clozapine-*n*-oxide. **(b)** Representative fluorescent image of hM4Di-mCherry expression in mOFC cell bodies and immunofluorescent image of mOFC axons and terminals in the BLA near the implanted guide cannula. **(c)** Schematic representation of hM4Di-mCherry expression in mOFC and axonal expression and injector tip placements in BLA for all subjects. **(d)** Lever-press rate (Lever presses/min) during the PIT test averaged across levers during the preCS baseline periods and during the CS periods on the lever earning the same outcome as the presented CS and the alternate lever (Different). Data are averaged across trials and CSs. **(e)** Rate of entries into the food-delivery port (Food-port entries/min) during the PIT test for the preCS baseline periods and during CS presentation (averaged across trials and CSs). **(f)** Average lever-press rate during the instrumental choice phase of the outcome-specific devaluation test. Presses separated for those on the action that in training earned the valued v. devalued (prefed) reward. **(g)** Rate of entries into the food-delivery port during the outcome-specific devaluation test, averaged across trials, during the preCS baseline period, and during presentation of the CS predicting the valued and devalued (prefed) rewards. Lines represent individual subjects. \**P* < 0.05, \*\**P* < 0.01, \*\*\**P* < 0.001.

Rats were food-deprived and given Pavlovian and instrumental conditioning using procedures that engender the encoding of rich, sensory-specific stimulus-outcome and action-outcome memories (Ostlund and Balleine, 2008; Malvaez et al., 2015; Lichtenberg and Wassum, 2016; Lichtenberg et al., 2017; Sias et al., 2021). For Pavlovian conditioning, each of 2, 2-min auditory conditional stimuli (CSs; white noise and tone) were associated with intermittent delivery of 1 of 2 distinct food rewards (sucrose solution or food pellets; e.g., white noise-sucrose/tone-pellet). Thus, each cue sets the ‘state’ in which a specific reward can be expected. During each session, each cue was presented 8 times in pseudorandom order (variable average 3-min intertrial interval) for 2 min, during which its associated reward was delivered on average every 30 s. Evidence of simple Pavlovian conditioning was detected in the goal-approach response (entries into the food-delivery port) to the cue before reward delivery (Baseline: 8.23 ± 0.89 s.e.m. entries/min; CS probe period: 18.72 ± 1.58; *t*_10_ = 12.57, *P* < 0.0001). Rats then received instrumental conditioning to learn that two different actions (left or right lever press) each earned one of the two food rewards (e.g., left press→sucrose/right press→pellets) ultimately on a random-ratio 20 schedule of reinforcement, thus establishing 2 action-outcome memories. Rats reached a final average press rate of 35.18 ± 3.13 presses/min.

We first asked whether activity in mOFC→BLA projections is necessary for the influence of stimulus-outcome and action-outcome memories over decision making using the outcome-specific Pavlovian-to-instrumental (PIT) test. Rats received two PIT tests, counterbalanced for order, one following intra-BLA infusion of CNO to inactivate hM4Di-expressing mOFC axons and terminals in the BLA and one following infusion of aCSF vehicle. We chose a vehicle-infused control to provide a within-subject comparison and based on evidence that CNO when infused at this dose into the BLA has no effect on the expression of PIT, similar reward-related behaviors, BLA activity, or OFC terminal activity in the BLA in the absence of the hM4Di transgene (Lichtenberg et al., 2017; Malvaez et al., 2019). During each test, both levers were continuously present, but lever pressing was not rewarded. Each CS was presented 4 times (also without accompanying reward), with intervening CS-free baseline periods, to assess its influence on action performance and selection in the novel choice scenario. The test was unrewarded to force subjects to rely on their memory of the predicted rewards. Because the cues are never associated with the instrumental actions, this test assesses the ability to, upon cue presentation, represent the specific predicted reward and use this to motivate choice of the action known to earn the same unique reward (Kruse et al., 1983; Colwill and Motzkin, 1994; Gilroy et al., 2014; Corbit and Balleine, 2016). If subjects are able use both the stimulus-outcome and action-outcome memories to enable accurate reward representation, then cue presentation should cause them to increase their lever presses selectively on the action earning the *same* outcome as predicted by that cue. This is consistent with the interpretation that rats use the cues to know which reward is predicted and, thus, infer which action would be most advantageous. Rats showed this outcome-specific PIT effect in the control condition (Figure 2d). Cue presentation increased presses selectively on the lever that, in training, earned the same outcome as the presented cue, relative to the lever that earned the different outcome. Conversely, the influence of the cues over lever-press choice was greatly attenuated following intra-BLA CNO infusion (Figure 2d; Drug × CS/Lever: *F*(1.4,14) = 4.20, *P* = 0.049; Drug (Vehicle v. CNO): *F*(_1,10_) = 9.51, *P* = 0.01; CS/Lever (preCS, CS-Same v. CS-Different): *F*(_1.5,14.7_) = 10.87, *P* = 0.002). This effect was selective to cue-influenced decision making. Inactivation of mOFC→BLA projections neither affected baseline lever-pressing activity (*P* > 0.9999, Bonferroni-corrected post-hoc comparison), nor conditional approach responses to the shared food-delivery port during the cues (Figure 2e; CS (Baseline v. CS): *F*(_1,10_) = 71.66, *P* < 0.0001; Drug: *F*(_1,10_) = 4.14, *P* = 0.07; Drug × CS: *F*(_1,10_) = 1.70, *P* = 0.22). Thus, activity in mOFC→BLA projections is not needed to support general reward pursuit activity or conditional approach responses but is necessary to use appetitive cues to know which reward is predicted and, thus, bias decision making.

PIT requires that the subjects both know which outcome is predicted by the presented cue and which action earns that same outcome. Thus, the disrupted expression of PIT by inactivation of mOFC→BLA projections could reflect a function for this pathway in stimulus-outcome memory, action-outcome memory, or both. To arbitrate between these possibilities, we retrained rats on both the Pavlovian and instrumental contingencies and gave them a set of outcome-specific devaluation tests. This also provided an opportunity to assess the contribution of mOFC→BLA projection activity to the ability to use the value of an anticipated reward to guide adaptive behavior. Prior to each test, one of the food rewards was devalued by sensory-specific satiety (1 hr prefeeding). No manipulation was made during this prefeeding to allow devaluation learning to proceed undisrupted. Immediately following the prefeeding, rats received an infusion of vehicle or CNO into the BLA and were then given a two-phase test. In the first phase, rats received 5-min access to both levers in a choice. Lever pressing was not reinforced to force subjects to rely on their action-outcome memories. If rats can use their action-outcome memories to represent the expected outcome of each action and its value, then they should select the action that earns the valued reward, downshifting responding on the action that earns the devalued reward. Rats showed this sensitivity of instrumental choice to outcome-selective devaluation in the control condition and when mOFC→BLA projections were inactivated (Figure 2f; Lever (Valued v. Devalued): *F*(_1,9_) = 15.32, *P* = 0.004; Drug: *F*(_1,9_) = 0.20, *P* = 0.67; Drug × Lever: *F*(_1,9_) = 0.47, *P* = 0.51). Thus, activity in mOFC→BLA projections is not necessary to retrieve action-outcome memories or use them to guide choice behavior. In the second phase, the levers were retracted and each cue was presented twice in alternating order. Again, the subjects were forced to rely on their memory, in this case their stimulus-outcome memories, because the cues were not accompanied by their associated reward. If rats can use the cues to know which reward is predicted and how valuable that reward currently is, then they should be able to infer that it is advantageous to respond to the cue signaling the valued outcome by checking the food-delivery port, but not advantageous to respond during the cue signaling the devalued outcome. This was apparent in the control condition. Following intra-BLA vehicle infusion, rats showed robust conditional food-port approach responses to the cue signaling the valued reward but attenuated responses to the cue signaling the devalued reward. Inactivation of mOFC→BLA projections prevented subjects from adapting their conditional goal-approach responses based on the value of the predicted reward (Figure 2g; Drug × CS: *F*(_1.3,11.9_) = 8.07, *P* = 0.01; Drug: *F*(_1,9_) = 1.32, *P* = 0.28; CS (Valued v. Devalued): *F*(_1.5,13.9_) = 20.47, *P* = 0.0001). Thus, activity in mOFC→BLA projections is necessary for adaptive responses to a cue based on the current value of the predicted reward. Together with the PIT results, this suggests that mOFC→BLA projection activity mediates the use of environmental cues to represent which specific reward is predicted and the desirability of that reward, both of which are critical for adaptive decision making.

### The BLA*→*mOFC pathway mediates adaptive cue responses based on the value of the predicted reward

We next asked whether BLA projections back to the mOFC are similarly involved appetitive behavior (Figure 3a; *N* = 9). The experiment was identical, except this time we expressed the inhibitory designer receptor hM4Di in the BLA and placed guide cannula bilaterally over the mOFC near hM4Di-expressing BLA axons and terminals (Figure 3b–c). This allowed later CNO (1 mM) infusion to inactivate BLA axons and terminals in the mOFC, as we have validated previously (Lichtenberg et al., 2017). We have also previously shown that in the absence of the hM4Di transgene CNO infusion at this dose into the OFC does not affect OFC activity, BLA terminal activity in the OFC, or reward-related behavior (Lichtenberg et al., 2017), and thus used a within-subject, vehicle-infused control condition. Rats provided evidence of successful Pavlovian conditioning in their final average goal-approach responses to the cues (Baseline: 6.60 ± 0.69 s.e.m. entries/min; CS probe period: 14.64 ± 1.71; *t*_8_ = 6.56, *P* = 0.0002) and also acquired performance of the instrumental actions, reaching a final average press rate of 39.70 ± 2.51 presses/min.

**Figure 3.**
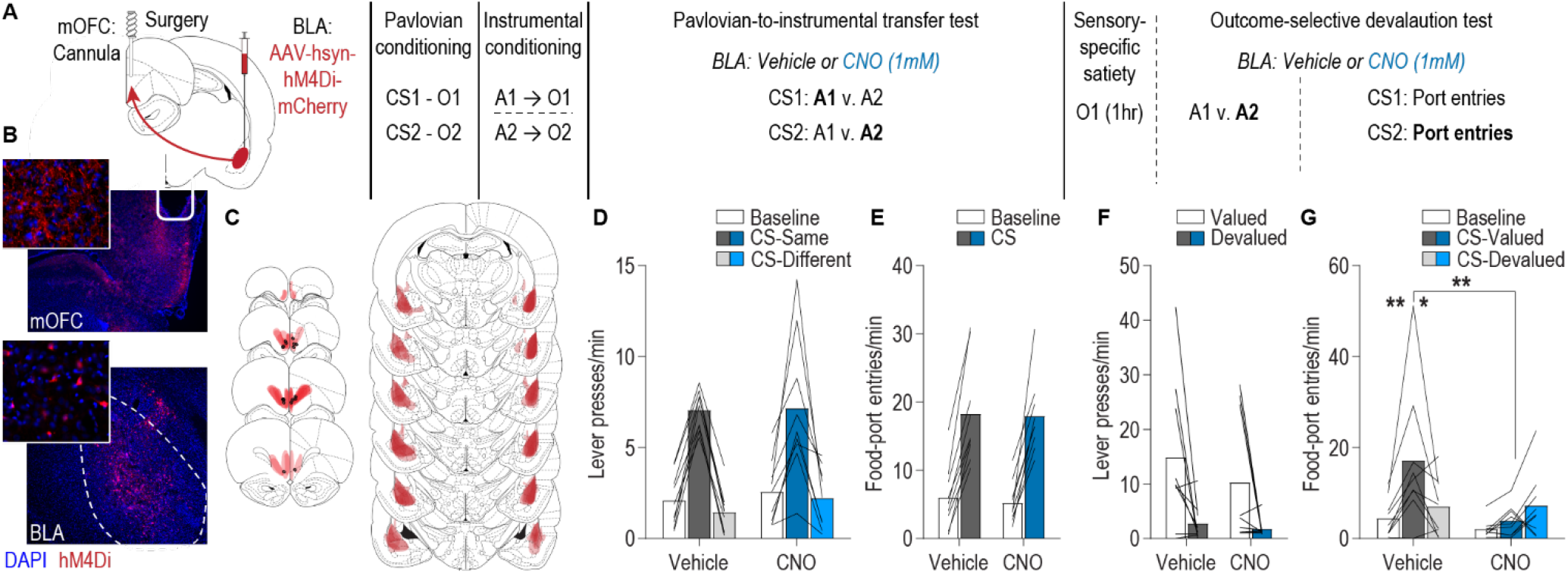
The BLA→mOFC pathway regulates adaptive cue responses based on predicted reward value. **(a)** Schematic of behavioral training, testing, and chemogenetic inactivation strategy. CS, conditional stimulus (white noise or tone); O, outcome (sucrose solution or food pellet); A, action (left or right lever press); CNO, clozapine-*n*-oxide. **(b)** Representative fluorescent image of hM4Di-mCherry expression in cell bodies of the BLA and immunofluorescent image of BLA axons and terminals in the mOFC near the implanted guide cannula. **(c)** Schematic representation of hM4Di-mCherry expression in BLA and axonal expression and injector tip placements in mOFC for all subjects. **(d)** Lever-press rate (Lever presses/min) during the PIT test averaged across levers during the preCS baseline periods and during the CS periods on the lever earning the same outcome as the presented CS and the alternate lever (Different). Data are averaged across trials and CSs. **(e)** Rate of entries into the food-delivery port (Food-port entries/min) during the PIT test for the preCS baseline periods and during CS presentation (averaged across trials and CSs). **(f)** Average lever-press rate during the instrumental phase of the outcome-specific devaluation test. Presses separated for those on the action that in training earned the valued v. devalued (prefed) reward. **(g)** Rate of entries into the food-delivery port during the outcome-specific devaluation test, averaged across trials, during the preCS baseline period, and during presentation of the CS predicting the valued and devalued (prefed) rewards. Lines represent individual subjects. \**P* < 0.05, \*\**P* < 0.01.

In contrast to the mOFC→BLA pathway, activity of BLA→mOFC projections was found to be unnecessary for the expression of PIT. Following intra-mOFC infusion of either vehicle or CNO, the reward-predictive cues were capable of biasing lever-pressing choice towards the lever that, during training, earned the same outcome as the presented cue relative to the lever that earned the different outcome (Figure 3d; CS/Lever: *F*(_1.5,12_) = 63.71, *P* < 0.0001; Drug: *F*(_1,8_) = 0.26, *P* = 0.62; Drug × CS/Lever: *F*(_1.2,9.5_) = 0.22, *P* = 0.69). During this PIT test, the conditional goal-approach response was also similar between groups (Figure 3e; CS: *F*(_1,8_) = 83.04, *P* < 0.0001; Drug: *F*(_1,8_) = 0.04, *P* = 0.86; Drug × CS: *F*(_1,8_) = 0.06, *P* = 0.81). Thus, activity in BLA→mOFC projections is neither needed to retrieve action-outcome or stimulus-outcome memories, nor to use this information to generate representations of the specific predicted reward to influence decision making.

Activity in BLA→mOFC projections was, however, critical to adapt approach responses to the cues based on the value of the predicted reward. During the outcome-specific devaluation test, in both the intra-mOFC vehicle and CNO conditions rats showed sensitivity of their instrumental choice behavior to outcome-specific devaluation, downshifting responding on the action that, in training, earned the devalued reward (Figure 3f; Lever: *F*(_1,8_) = 20.60, *P* = 0.002; Drug: *F*(_1,8_) = 0.57, *P* = 0.47; Drug × Lever: *F*(_1,8_) = 0.22, *P* = 0.65). This confirmed that BLA→mOFC projection activity is not needed to retrieve action-outcome memories and further demonstrated that these projections are not needed for general value-based decision making. However, inactivation of BLA→mOFC projections did prevent subjects from adapting their Pavlovian conditional food-port approaches responses based on the value of the predicted reward. Following intra-mOFC vehicle infusion, rats showed robust conditional food-port approach responses to the cue signaling the valued reward and attenuated responses to the cue signaling the devalued reward. This sensitivity of the conditional goal-approach response to devaluation was abolished following intra-mOFC CNO infusion (Figure 3g; Drug × CS: *F*(_2,16_) = 5.01, *P* = 0.02; Drug: *F*(_1,8_) = 2.98, *P* = 0.12; CS (Valued v. Devalued): *F*(_2,16_) = 9.86, *P* = 0.002). Thus, based on the PIT results, activity in the BLA→mOFC pathway is not needed to use the cues to know the identity of predicted reward, but the devaluation results indicate it is required to use the current value of that reward to infer how advantageous it would be to respond to the cue.

## DISCUSSION

Here we interrogated the function of mOFC-BLA circuitry in the prospective considerations that underlie adaptive reward-related behavior. Using anatomical tracing, we confirmed the existence of bidirectional, direct pathways between mOFC and BLA and found that BLA projections to the mOFC and lOFC are largely distinct. Using pathway-specific chemogenetic inhibition, we found that mOFC→BLA projection activity is critical for using stimulus-outcome memories to guide decision making based on which specific reward is expected and adaptive cue responses based on the desirability of that predicted outcome. BLA→mOFC projection activity, however, is only needed to adapt cue responses based on the desirability of the predicted reward.

We found that mOFC→BLA pathway activity is critical for using predictive environmental cues to know which specific reward is predicted and the current value of that option. mOFC neuronal activity can represent a cue-reward memory (Namboodiri et al., 2019) and is necessary, across species, for the use of such memories to inform adaptive behavior (Noonan et al., 2010; Bradfield et al., 2015; Noonan et al., 2017; Bradfield et al., 2018). Our data indicate this function is, at least in part, achieved via direct projections to the BLA. We have previously found that mOFC→BLA pathway activity mediates the retrieval of the incentive value of an expected food reward to ensure its adaptive pursuit (Malvaez et al., 2019). Incentive value is another form of state-dependent reward memory (food has high value when hungry, but low when sated). Thus, mOFC→BLA projections may be responsible for using the current state, defined both by external and internal physiological cues, to make a judgement about how advantageous a certain course of action might be.

This function of mOFC→BLA projections is distinct from that of lOFC→BLA projections. Using the same procedures, we previously found that lOFC→BLA projections are not needed for reward cues to bias choice towards the predicted reward (Lichtenberg et al., 2017). They are also unnecessary for retrieving the incentive value of an expected reward (Malvaez et al., 2019). Rather, lOFC→BLA pathway activity drives the learning of these state-dependent reward memories (Malvaez et al., 2019; Sias et al., 2021), a function that requires the BLA itself (Wassum et al., 2009; Wassum et al., 2011; Parkes and Balleine, 2013; Wassum et al., 2016), but not mOFC→BLA projections (Malvaez et al., 2019). Thus, the lOFC→BLA pathway mediates the formation of state-dependent reward memories and the mOFC→BLA pathway facilitates the use of this information to guide adaptive reward-related behavior. Interestingly, however, under more dynamic decision scenarios the function of individual components of the OFC-BLA circuit can appear more overlapping. For example, during reversal learning subjects must learn, integrate, and use information about reward availability and option value, and this form of decision making is influenced by lesion of lOFC→BLA projections (Groman et al., 2019).

The BLA→mOFC pathway was found to mediate adaptive responses to cues based on the desirability of the predicted reward. This is consistent with prior evidence that the BLA is needed for sensitivity of Pavlovian responses to outcome-specific devaluation (Hatfield et al., 1996; Johnson et al., 2009) and evidence that BLA neuronal responses to reward-predictive cues can reflect the value of the predicted reward (Schoenbaum et al., 1998; Saddoris et al., 2005; Paton et al., 2006; Belova et al., 2007; Belova et al., 2008). The data here reveal this function to be achieved, at least in part, via projections to the mOFC. BLA→mOFC projection activity was not necessary for reward cues to bias choice towards the predicted reward. Expression of such outcome-selective PIT depends on the sensory-specific identity of the predicted reward, but not its value (Holland, 2004). Thus, BLA→mOFC pathway activity mediates cue-driven behaviors based on a representation of the predicted reward's value, but not its identity. Surprisingly, this indicates that a reward's identity can be decoupled from its value; that one can represent which specific reward is predicted, but not the current value of that reward.

The function of the BLA→mOFC pathway identified here is distinct from that previously identified for the BLA→lOFC pathway. Using the same procedures, we previously found that activity in the anatomically distinct BLA→lOFC pathway is necessary for reward cues to both bias action choice towards the predicted reward and to adapt conditional approach responses based on the value of the predicted reward. Thus, the BLA→lOFC pathway allows one to use cues to know which specific reward is predicted, whereas BLA→mOFC pathway activity promotes behavior based on the current desirability of that predicted reward. Whether BLA→lOFC function in value is secondary to representing reward identity (if you do not know which reward is predicted, then you cannot represent its value) is a critical open question.

Neither mOFC→BLA, nor BLA→mOFC projections were necessary for general conditional goal-approach responses, consistent with evidence from BLA or mOFC lesions (Hatfield et al., 1996; Everitt et al., 2000; Parkinson et al., 2000; Corbit and Balleine, 2005; Bradfield et al., 2015; Malvaez et al., 2015; Bradfield et al., 2018; Morse et al., 2020). This well-learned cue response does not require a representation of the specific predicted reward or on-the-fly use of this information for prospective consideration of the most advantageous option. Instead, it can rely on a previously learned policy. Thus, mOFC-BLA circuitry is not needed for cue-responses generally but is needed when one must use stimulus-outcome memories to infer what to do based on which outcomes are expected and their current desirability.

The mOFC-BLA circuit was also not needed for rats to adapt their instrumental choice behavior based on the value of the predicted reward. Rather, it is selectively needed for cues to inform the considerations guiding adaptive appetitive behavior. Both the BLA and mOFC are needed for the sensitivity of instrumental choice to devaluation (Balleine et al., 2003; Ostlund and Balleine, 2008; Johnson et al., 2009; Bradfield et al., 2015; Gourley et al., 2016; Bradfield et al., 2018). This function may, therefore, be achieved via alternate pathways, perhaps those to the striatum (Hoover and Vertes, 2011; Corbit et al., 2013; van Holstein et al., 2020), a region heavily implicated in action-outcome memory (Malvaez et al., 2018b; Malvaez and Wassum, 2018; Malvaez, 2019).

Collectively these data reveal the mOFC-BLA circuit as critical for the cue-dependent reward outcome expectations that influence decision making and adaptive cue responses. mOFC neuronal activity can represent partially unobservable states (Lopatina et al., 2017; Elliott Wimmer and Büchel, 2019) and is needed when such states must be used to make adaptive choices (Bradfield et al., 2015; Bradfield et al., 2018). Activity in the mOFC→BLA pathway may, therefore, be involved in signaling the state predicted by the reward cues. The BLA pathways to the lOFC and mOFC might then facilitate the use of this information to guide adaptive behavior, with BLA→lOFC projections facilitating detailed, sensory-specific representations of expected rewards (Lichtenberg et al., 2017) and BLA→mOFC projections facilitating the more general representations needed to compute option desirability. Indeed, lOFC neuronal activity can encode high-dimensional representations and the identity of predicted rewards (Klein-Flügge et al., 2013; McDannald et al., 2014; Rudebeck and Murray, 2014; Wilson et al., 2014; Howard et al., 2015; Rich and Wallis, 2016; Suzuki et al., 2017; Rudebeck and Rich, 2018), whereas mOFC represents more general information about expected events that is used to make decisions based on value estimations or comparisons (Pritchard et al., 2005; Padoa-Schioppa and Assad, 2006; Plassmann et al., 2010; Kennerley et al., 2011; Levy and Glimcher, 2011; Rudebeck and Murray, 2011; Burton et al., 2014; Lopatina et al., 2016; Lopatina et al., 2017; Suzuki et al., 2017). The precise information content conveyed by each component of the OFC-BLA circuit and how it is used in the receiving structure is a critical question for follow-up investigation. Because the staged scenarios used here allow isolation of psychological processes that typically co-occur and interact to influence decision making, such activity analysis will also be important for understanding how the functions identified here relate to more dynamic and complex decision-making scenarios. Another essential question is whether mOFC-BLA circuit function is similar in females, who do show similar performance in the tasks used here and require the BLA and mOFC for their performance (Ostlund and Balleine, 2008; Bradfield et al., 2018).

An inability to use reward cues to inform prospective considerations of which specific rewards can be expected and their current desirability can lead to maladaptive choices. Maladaptive choices could also arise if one is able to know what rewards are predicted, but not to consider their current value, as we showed here is neurobiologically possible. This is characteristic of the cognitive symptoms underlying many psychiatric diseases, including substance use disorder (Kalivas and Volkow, 2005; Verdejo-Garcia et al., 2018). Moreover, both the mOFC and BLA, as well as their connectivity, can be dysfunctional in substance use disorder and other psychiatric illnesses marked by disrupted appetitive decision making (Tanabe et al., 2009; Goldstein and Volkow, 2011; Hahn et al., 2011; Linke et al., 2012; Sladky et al., 2015; Shields and Gremel, 2020; Xie et al., 2021). Thus, these data may aid our understanding and treatment of these conditions.

## DATA AND CODE AVAILABILITY

All data supporting the findings of this study are available from the corresponding author upon request.

## ADDITIONAL INFORMATION

### Funding

| **Funder** | **Grant reference number** | **Author** |
| --- | --- | --- |
| National Institute of Drug Abuse | DA035443 | Kate M. Wassum |
| National Institute of Drug Abuse | F31DA044734 | Nina Lichtenberg |
| National Institute of Drug Abuse | T32DA024635 | Nina Lichtenberg (PI: London) |
| National Institute of Drug Abuse | T32DA024635 | Alexander Lamparelli (PI: London) |

### Author contributions

NTL and KMW designed the research, analyzed and interpreted the data, and wrote the manuscript. NTL conducted the tracing and chemogenetic experiments. ZTP contributed to the analysis of the tracing data. LS conducted the mOFC→BLA chemogenetic experiments, with assistance from VYG. ACL contributed to the devaluation experiments.

### Ethics

#### Animal experimentation

All procedures were in accordance with the US National Institutes of Health (NIH) Guide for the Care and Use of Laboratory Animals and were approved by the UCLA Institutional Animal Care and Use Committee (protocol 2012-021).

## Acknowledgements

We would also like to acknowledge the very helpful feedback from Drs. Alicia Izquierdo, Melissa Sharpe, and Melissa Malvaez on this data. Lastly, we would like to acknowledge the generous infrastructure support from the Staglin Center for Behavior and Brain Sciences.

